# Systematic mutagenesis reveals critical effector functions in the assembly and dueling of the H1-T6SS in *Pseudomonas aeruginosa*

**DOI:** 10.1101/2022.09.14.507963

**Authors:** Li-Li Wu, Tong-Tong Pei, Shuangquan Yan, Ming-Xuan Tang, Li-Wen Wu, Hao Li, Xiaoye Liang, Shuyang Sun, Tao Dong

## Abstract

*Pseudomonas aeruginosa* is an important human pathogen that can cause severe wound and lung infections. It employs the type VI secretion system (H1-T6SS) as a molecular weapon to carry out a unique dueling response to deliver toxic effectors to neighboring sister cells or other microbes after sensing an external attack. However, the underlying mechanism for such dueling is not fully understood. Here, we examined the role of all H1-T6SS effectors and VgrG proteins in assembly and signal sensing by ectopic expression, combinatorial deletion and point mutations, and imaging analyses. Expression of effectors targeting the cell wall and membrane resulted in increased H1-T6SS assembly. Deletion of individual effector and *vgrG* genes had minor-to-moderate effects on H1-T6SS assembly and dueling activities. The dueling response was detectable in the *P. aeruginosa* mutant lacking all H1-T6SS effector activities. In addition, double deletions of *vgrG1a* with either *vgrG1b* or *vgrG1c* and double deletions of effector genes *tse5* and *tse6* severely reduced T6SS assembly and dueling activities, suggesting their critical role in T6SS assembly. Collectively, these data highlight the diverse roles of effectors in not only dictating antibacterial functions but also their differential contributions to the assembly of the complex H1-T6SS apparatus.

## Introduction

The gram-negative *Pseudomonas aeruginosa* is one of the ESKAPE pathogens of high clinical relevance owing to its acquisition of a plethora of genes encoding antibiotic resistance and virulence factors ^1–3^. To develop effective treatment strategies, it is important to understand how *P. aeruginosa* survives in polymicrobial communities that it encounters during infection. To outcompete other bacteria, *P. aeruginosa* employs the type VI secretion system (T6SS) as a powerful weapon to translocate effector proteins to the extracellular environment or directly into target cells.

The T6SS is a contractile injection device composed of an outer contractile sheath (TssB/TssC), an inner needle (Hcp) and a puncturing spike (VgrG and PAAR) ^4–7^. The outer sheath is anchored to the cell envelope through the baseplate complex (TssEFGK) which acts as a platform for the assembly of the contractile tail and connects it to the membrane complex(TssJLM) ^8–11^. Contraction of the sheath propels the needle and the spike complex out of the cells to penetrate neighboring cell membranes, releasing VgrG, PAAR, Hcp subunits and effectors with different functions into target cells ^12–14^. Hcp, VgrG or PAAR proteins can carry additional C-terminal domains as specialized effectors or non-covalently interact with cargo effectors ^15–20^. TssA is essential for the assembly of T6SS, acting as chaperones assisting sheath-tube polymerization ^11,21–24^. The AAA+ ATPase ClpV can recognize and disassemble the contracted sheath complex ^25–27^.

The T6SS has been found in 25% of gram-negative bacteria ^28–30^, and there are three separate T6SS gene clusters (H1-, H2-, and H3-T6SS) encoded in the *P. aeruginosa* model strain PAO1 genome ^30^. The H1-T6SS mainly targets bacteria while the H2- and H3-T6SSs mediate bacterial competition, metal acquisition, and interaction with host cells ^18,31–33^. For the eight known effectors (Tse1-8) of H1-T6SS, they can be categorized into two groups: periplasmic-targeting effectors (Tse1, and Tse3-5) and cytosolic-targeting effectors (Tse2, and Tse6-8) ^16,31,34–38^. These effectors may function synergistically or exert conditional toxicities against different competitors ^36^. Tse1 and Tse3 hydrolyze peptidoglycan, acting as an amidase and a muramidase, respectively ^35^. Tse4 is a pore-forming toxin that disrupts proton motive force (PMF) ^36^ while Tse5 is toxic in the periplasm with unknown mechanism ^31^. For cytosolic toxins, Tse2 is a predicted NAD-dependent ADP-ribosyltransferase, Tse6 degrades NAD^+^ and NADP^+^, Tse7 is a DNase with a Tox-GHH2 domain, and Tse8 targets the transamidosome complex in bacteria ^16,37–39^. Several substrates of H2-T6SS have also been discovered, including phospholipase effectors PldA and PldB, TOX-REase-5 domain-containing effector TseT, molybdate-binding protein ModA, and PA2066 ^18,40–43^. The H3-T6SS secreted effectors include TseF that facilitates metal ion acquisition ^33^ and PA0256, a putative antibacterial effector that binds Hcp3 ^42^.

The expression and assembly of T6SS in *P. aeruginosa* is tightly regulated at the transcriptional and posttranslational levels. H1-T6SS and H2-T6SS are up-regulated in a Δ*retS* mutant ^18,30^. The deletion of the *retS* gene activates the GacS/GacA two-component system and GacA drives the expression of two small noncoding RNAs, RsmY and RsmZ, which sequester the T6SS-inhibitory small noncoding RNA RsmA ^44–46^. The H1-T6SS is normally not actively assembled unless provoked by a T6SS-active competitor or other external membrane-perturbing signals including EDTA, extracellular DNA, and polymyxin B ^47–50^. This response is termed dueling/tit-for-tat and controlled at the post-translational level by the threonine phosphorylation pathway (TPP) consisting of the TagQRST sensor and the PpkA-PppA kinase-phosphatase pair. PpkA phosphorylates an H1-T6SS essential protein Fha1 that leads to H1-T6SS activation while the PppA counteracts with this process ^49,51– 53^. When *P. aeruginosa* is co-incubated with *Vibrio cholerae, P. aeruginosa* could differentiate T6SS^+^ from T6SS^−^ *V. cholerae* in a tripartite mix by primarily sensing the damage caused by a delivered *V. cholerae* lipase effector TseL, while the other effectors or the physical puncture of *V. cholerae* T6SS induced much weaker responses ^49,54^. Removal of T6SS effectors in *Agrobacterium tumefaciens* abolished tit-for-tat response in *P. aeruginosa* ^55^. Endogenous membrane stress triggered by deletion of *bamA, tolB*, and *lptD* genes involved in the biogenesis of outer membranes, could also induce T6SS activation in *P. aeruginosa* ^56^. However, the molecular details for T6SS dueling between *P. aeruginosa* cells remains elusive.

Here we focus on elucidating the effects of H1-T6SS effectors on T6SS assembly and competition by systematically constructed all effector mutants and expression vectors and determined their effects on T6SS assembly, dueling, and competition in *P. aeruginosa*. Our results show that T6SS dueling is induced by H1-T6SS but not H2-T6SS or H3-T6SS. Although none of the individual effector was required for dueling and assembly, periplasmic expression of the peptidoglycan-targeting Tse1 induced the T6SS substantially. VgrG1a, VgrG1b and VgrG1c also contribute to the T6SS dueling, and their corresponding effectors, Tse5-Tse7, are critical for T6SS assembly as structural components. The *8eff*_*c*_ mutant, lacking all effector activities, still exhibited dueling between sister cells and it could deliver a Cre-recombinase into a T6SS^+^ *V. cholerae* non-toxic reporter to induce recombination. These data may suggest the dueling response is highly sensitive and can sense both effector-elicited damages and the T6SS physical penetration.

## Results

### Dueling is restricted to H1-T6SS pairs

We first examined whether dueling of the H1-T6SS is modulated by the other two separate T6SS clusters in *P. aeruginosa*. Because the deletion of *retS* can increase the expression of both H1-T6SS and H2-T6SS ^18,30,48^, we tested whether H2- and H3-T6SSs contribute to the T6SS dueling in the *ΔretS* mutant background. We constructed single T6SS active mutants, H1-, H2-, and H3-T6SS^+^ strains, by inactivating the other two T6SSs in the *ΔretS* mutant, respectively. We also labeled each corresponding TssB protein with sfGFP (Superfolder GFP) to monitor T6SS assemblies by time-lapse fluorescence microscopy. We found that paired TssB1-sfGFP could be observed in the H1-T6SS^+^ cells (Fig. 1A-1C and Video S1). Only unpaired discrete foci of TssB2-sfGFP signals were observed in the H2-T6SS^+^ cells (Fig. 1A-1C and Video S2). No TssB3-sfGFP signals were observed in the H3-T6SS^+^ mutants (Fig. 1A-1C and Video S3) suggesting H3-T6SS is not active under the tested conditions. To determine whether H2-T6SS can activate the assembly of H1-T6SS, we mixed sfGFP-labeled H2-T6SS^+^ with mCherry2-labeled H1-T6SS^+^ (Δ*tssB2*, Δ*tssB3*, TssB1-mCherry2) at an equal ratio (Fig. 1D and Video S4). No H1-T6SS/H2-T6SS dueling pair were detected (Video S4), suggesting that H2-T6SS does not stimulate the assembly of H1-T6SS. These results confirm that dueling is specific to the H1-T6SS.

**Fig. 1.**
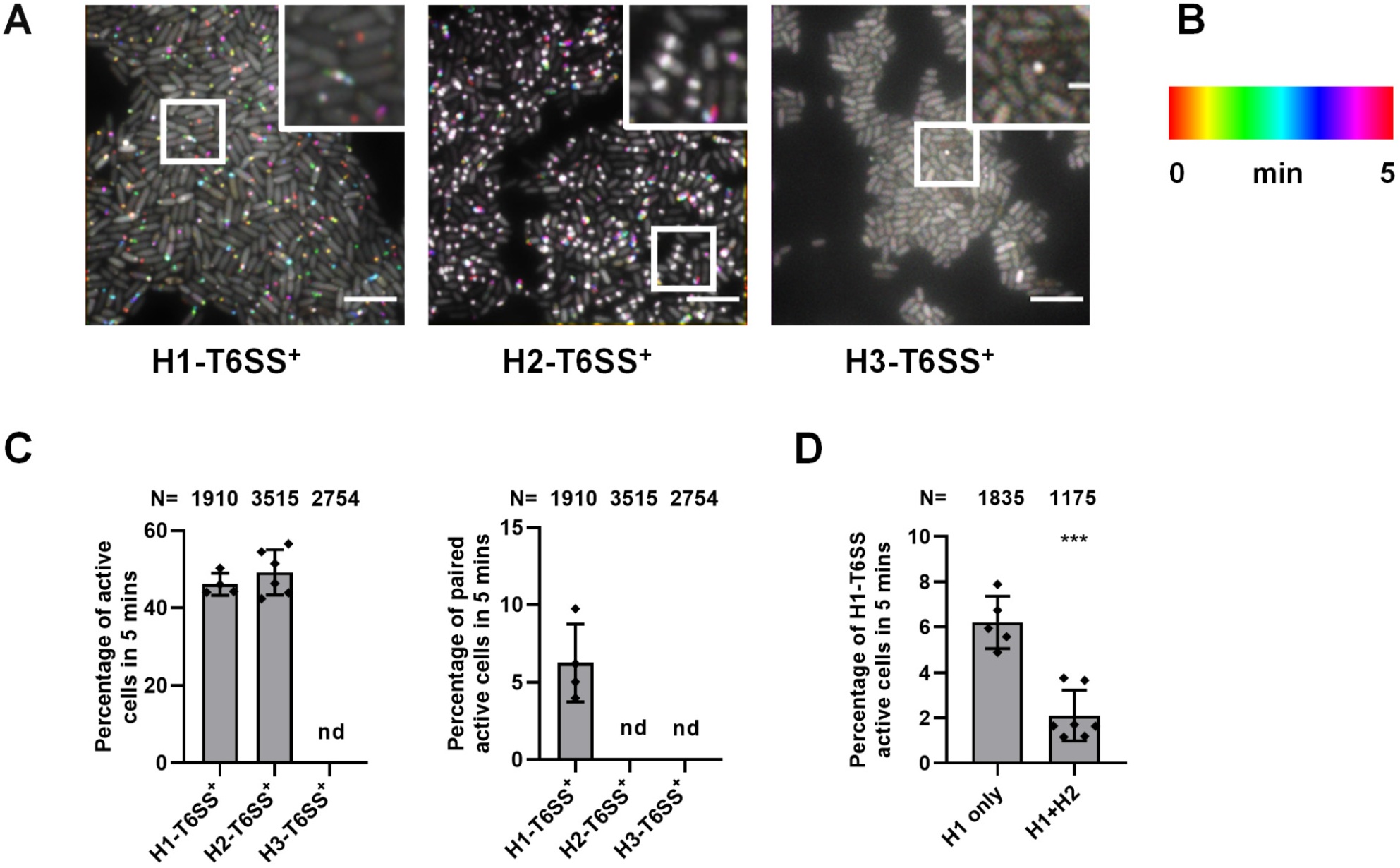
Dueling is restricted to H1-T6SS pairs. **A**, Time-lapse imaging of TssB1-sfGFP, TssB2-sfGFP, and TssB3-sfGFP signal captured every 10 s for 5 min and temporally color-coded. H1-T6SS^+^ mutant cells were collected at OD_600_∼1, H2-T6SS^+^ mutant cells were collected at OD_600_∼2 and H3-T6SS^+^ mutant cells were collected at OD_600_∼3. A representative image of 30 μm × 30 μm field of cells with a 2 × magnified 5 μm × 5 μm inset (marked with a box) of a selected region is shown. Scale bar in A is 5 μm for the large field of view and 1 μm for the insets view. Strain genotypes are indicated at the bottom. H1-T6SS^+^: Δ*retS, ΔtssB2, ΔtssB3*, TssB1-sfGFP; H2-T6SS^+^: Δ*retS*, Δ*tssB1, ΔtssB3*, TssB2-sfGFP; H3-T6SS^+^: Δ*retS, ΔtssB1, ΔtssB2*, TssB3-sfGFP. **B**, Color scale used to temporally color code the sfGFP signal. **C**, Quantification of active cells forming sheaths (left) and paired active cells (right) counted in *P. aeruginosa ΔretS* with only one T6SS mutants. Error bars indicate the mean ± standard deviation of at least three different biological duplicates with at least four fields. nd, not detected. **D**, Quantification of active H1-T6SS when H1-T6SS^+^ (Δ*tssB2, ΔtssB3*, TssB1-mCherry2) cells were cultured without/with H2-T6SS^+^ cells (H1 only or H1+H2). Error bars indicate the mean ± standard deviation of at least two different biological duplicates with more than five fields. One-way ANOVA with Dunnett’s multiple comparisons test compared with H1-T6SS^+^ was used in C and a two-tailed Student’s *t*-test was used in D. N indicates the total number of cells counted for each strain per a 30 μm × 30 μm field in C and D. \*\*\**P* < 0.001.

### Inactivation of single H1-T6SS secreted effectors does not abolish T6SS dueling in *P. aeruginosa*

Because the lipase effector in *V. cholerae* has been shown to be the primary effector provoking H1-T6SS firing in *P. aeruginosa*, we next investigated whether specific effectors of the H1-T6SS play a similar role in stimulating T6SS dueling. We constructed eight single effector deletion mutants by knocking out each effector and measured T6SS dueling with microscopy. Although all effector mutants exhibited T6SS assembly and dueling pairs, the number of dueling pairs in the *tse1, tse4, tse6*, and *tse7* mutants was relatively reduced (Fig. 2A-2B and Fig. S1A). Therefore, unlike in *V. cholerae*, there is no specific sole effector responsible for H1-T6SS dueling in *P. aeruginosa*.

**Fig. 2.**
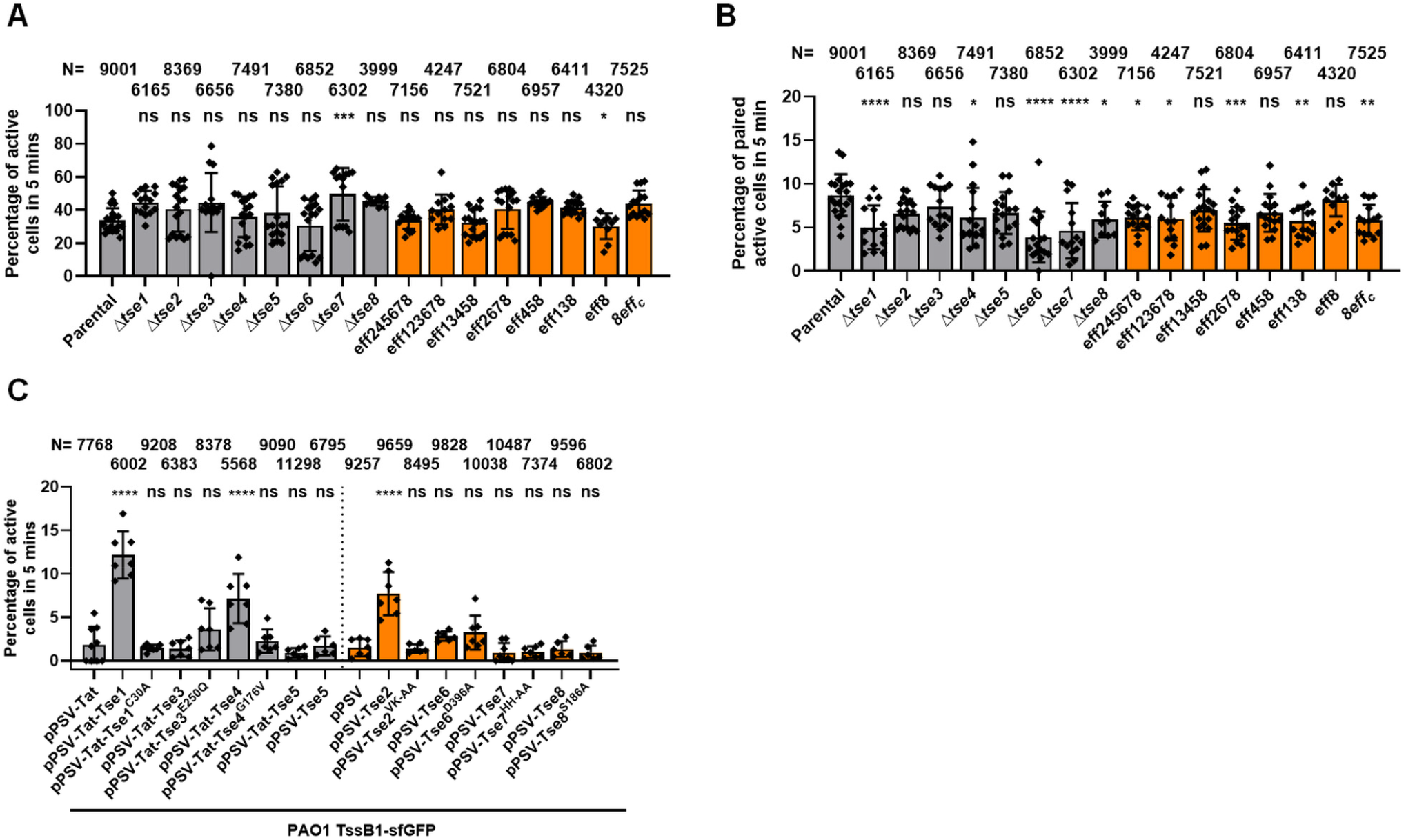
Tse effectors targeting cell membrane and peptidoglycan contribute to T6SS activation and dueling. **A-B**, Quantification of active cells forming sheaths (A) and paired active cells (B) counted in *P. aeruginosa* H1-T6SS^+^ with the deletion of *tse* genes individually or with combinatorial effector inactivation detected with per a 30 μm × 30 μm field. Parental: H1-T6SS^+^; eff: mutants with active effectors after combinatorial effector inactivation; *8eff*_*c*_: eight-effector-inactivated mutant. Time-lapse imaging figures are shown in Fig S1A and S1B. **C**, Quantification of active cells forming sheaths in *P. aeruginosa* TssB1-sfGFP strain expressing Tse effectors. Tse1, Tse1^C30A^, Tse3, Tse3^E250Q^, Tse4, Tse4^G176V^ and Tse5 were expressed in the periplasm (Tat-signal added). *P. aeruginosa* strains expressing indicated plasmids were detected per a 132 μm × 132 μm field. *P. aeruginosa* expressing pPSV-Tat or pPSV is used as control. Toxicity assay of Tse effectors in *E. coli* or *P. aeruginosa* TssB1-sfGFP strain is shown in Fig S2A-S2H and S3A-S3H, respectively. Error bars indicate the mean ± standard deviation of at least three different duplicates with more than ten fields in A and B or more than six fields in C. One-way ANOVA with Dunnett’s multiple comparisons test compared with the parental strain in A-B or with *P. aeruginosa* TssB1-sfGFP strain with pPSV or pPSV-Tat in C. \**P* < 0.05, \*\**P* < 0.01, \*\*\**P* < 0.001, \*\*\*\**P* < 0.0001; ns, not significant.

### Tse effectors targeting cell membrane and peptidoglycan contribute to T6SS activation and dueling

Given that all the H1-T6SS effector single deletion mutants still had dueling pairs, we next investigated whether dueling is dependent on specific toxicities conferred by multiple redundant effectors. Therefore, we grouped effectors based on their toxicities in the periplasm or the cytoplasm. Considering some effectors may also contribute to assembly, as previously reported in other species ^57,58^, we generated in-frame deletion mutation of Tse5, as the active site remains unclear ^31^, and catalytic mutations of the other seven effectors, resulting in Tse1^C30A^, Tse2^VK-AA^ (residues 109-110, valine, lysine, to alanine residues), Tse3^E250Q^, Tse4^G176V^, Tse6^D396A^, Tse7^HH-AA^ (residues 229-230, histidine, histidine, to alanine residues) and Tse8^S186A 16,35–38,59^. We constructed a series of select combinatorial effector inaction/deletion mutants in H1-T6SS^+^ (Δ*retS*, Δ*tssB2*, Δ*tssB3*, TssB1-sfGFP) background to determine if specific activities were required to elicit dueling, including a mutant lack peptidoglycan-damaging activities (eff245678) and a mutant lacking cell membrane-damaging effectors (eff123678). Interestingly, all effector mutants had similar levels of T6SS assembling activity comparable to the parental strain (Fig. 2A-2B and Fig. S1B). The number of dueling pairs decreased significantly when effectors targeting peptidoglycan or cell membrane (eff245678, eff123678 and eff2678) were inactivated (Fig. 2A-2B and Fig. S1B), suggesting that the peptidoglycan targeting Tse1 and Tse3, or cell membrane targeting Tse4 and Tse5 may contribute to the H1-T6SS dueling.

To compare with effector-inactivation effects, we also tested whether expression of H1-T6SS effectors could stimulate the H1-T6SS assembly using an IPTG-inducible expression vector pPSV37. The corresponding effector-inactive mutants were also tested as control. The toxicities of all constructs were examined in *E. coli* and the *P. aeruginosa* TssB1-sfGFP strain (Fig. S2A-S2H, Fig. S3A-S3H, and Fig.S4). Based on their known toxicities, Tse1, Tse3, Tse4, and Tse5 were expressed in the periplasm with the Tat (two-arginine translocation) secretion signal while Tse2, Tse6, Tse7, and Tse8 were expressed in the cytoplasm. Microscopy analysis showed that overexpression of peri-Tse1, Tse2 and peri-Tse4, but not the other constructs, stimulated the T6SS assembly significantly in wild-type *P. aeruginosa* compared with their catalytically inactive variants and the empty plasmid control (Fig. 2C and Fig. S5A-S5B). Cells overexpressing peri-Tse1 became spherical but the overexpression of other effectors in *P. aeruginosa* did not lead to obvious morphologic changes (Fig. S5A-S5B).

Considering that cognate immunity proteins in *P. aeruginosa* may neutralize the toxicity of effectors, we also expressed Tse effectors and tested their toxicity in their effector-immunity deletion mutants (Fig. S6A-S6G). Microscopy analysis showed that the expression of peri-Tse1 in the Δ*tse1*^*ei*^ exhibited the most T6SS activation (Fig. S7A, S7I). Cells expressing Tse2, peri-Tse4 and peri-Tse5 in their cognate effector-immunity mutants showed a moderate increase of T6SS assemblies (Fig. S7B, S7D, S7E, S7I), whereas the expression of peri-Tse3 and Tse7 showed no significant increase of T6SS assemblies compared with control strains (Fig. S7C, S7G, S7I). Interestingly, cells expressing Tse6 exhibited a decrease in T6SS activation while cells expressing Tse6^D396A^ showed a significant increase in T6SS assembly compared with their control strain (Fig. S7F, S7I). These results suggest that Tse1-mediated cell-wall damage plays a major role in the T6SS activation and dueling in *P. aeruginosa*.

### Tse1-mediated H1-T6SS activation is dependent on TagT

Because the TagQRST-PpkA-Fha1-PppA regulatory cascade can sense *V. cholerae* T6SS attack and the TseL effector activity at the post-translational level ^49,54^, we next tested whether Tse1-mediated cell-wall damage could trigger assembly of the H1-T6SS in the Δ*tagT* mutant. Expression of Tse1 in the periplasm resulted in dramatic changes in cell shapes and cell lysis, indicative of cell-wall damages, but did not lead to increased H1-T6SS assembly relative to the catalytically inactive Tse1^C30A^ mutant or the empty vector controls (Fig. S8A-S8C). These results suggest that the TagQRST pathway is required for sensing Tse1-mediated cell-wall damages.

### *P. aeruginosa* employs Tse1-Tse7 but not Tse8 to kill competing pathogens

Having built a panel of combinatorial effector mutants, we next tested effector-killing specificities against three T6SS-active pathogens, *V. cholerae* V52, *Aeromonas dhakensis* SSU, and an emerging pathogen *Enterobacter cloacae* B29. All except for the Tse8-only effector mutants exhibited wild-type-level killing against *V. cholerae* V52, *A. dhakensis* SSU and *E. cloacae* B29 (Fig. 3A-3C and Fig. S9A-S9C). These results suggest that these effectors display general toxicities against competing bacteria.

**Fig. 3.**
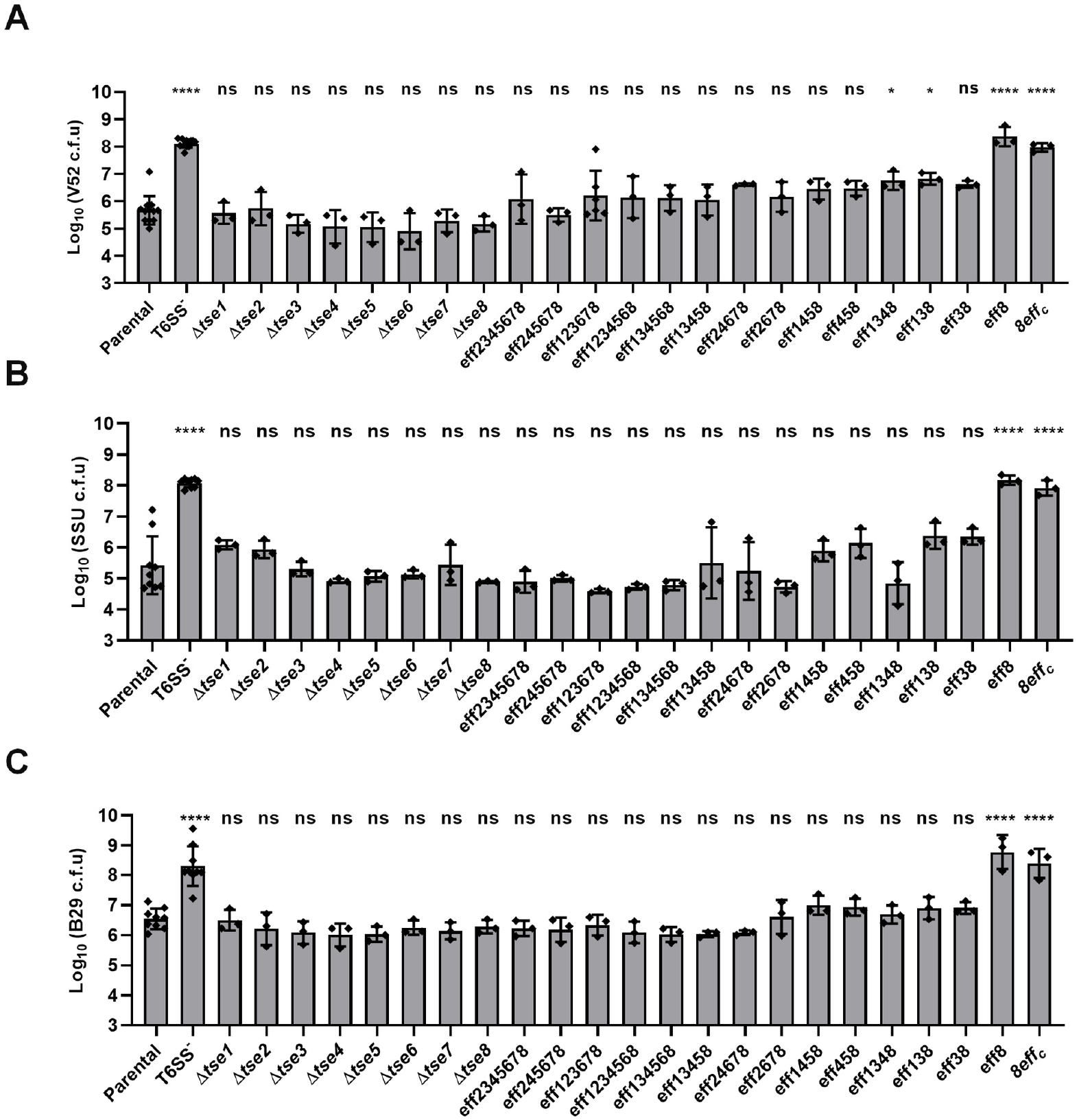
Tse1-Tse7 but not Tse8 exhibit killing ability against competing pathogens. **A-C**, Survival of *V. cholerae* V52 (A), *A. dhakensis* SSU (B) and *E. cloacae* B29 with pBBR1MCS-5 plasmid (C) after competition assay against PAO1 mutants. Genotypes of PAO1 were indicated at the bottom. Survival of PAO1 is shown in Fig S9A-S9C. Killer and prey cells were mixed at the ratio of 5: 1. Error bars indicate the mean ± standard deviation of at least three different biological duplicates. One-way ANOVA with Dunnett’s multiple comparisons test compared with the parental strain. \**P* < 0.05, \*\*\*\**P* < 0.0001; ns, not significant.

### Deletion of *vgrG* and the cognate effector genes reduces T6SS dueling

Because the VgrG-spike complex promotes the T6SS assembly ^5,60–62^, we next asked whether the three VgrG proteins (VgrG1a/b/c) contribute to T6SS dueling. Deletion of *vgrG1a-c* genes individually had little effect on T6SS assembly, confirming previous findings ^62,63^, but the T6SS dueling was reduced by half relative to the wild-type level (Fig. 4A-4B and Fig. S10A-S10D). Double deletion of *vgrG1a* with either *vgrG1b* or *vgrG1c* abolished T6SS assembly while the *vgrG1b*/*c* double deletion mutant maintained T6SS assembly (Fig. 4A-4C), suggesting that VgrG1a plays a key role in T6SS assembly but can be substituted by a heterotrimeric VgrG1b/c. These data also suggest that the delivered sister-cell VgrGs can facilitate T6SS assembly and dueling. Because each VgrG has its cognate effector (VgrG1a-Tse6, VgrG1b-Tse7, VgrG1c-Tse5) ^38,63,64^, we also examined the effect of combinatorial deletion of effector genes on the H1-T6SS. Double deletion of *tse5* and *tse6* severely impaired T6SS assembly, mimicking the effect of their corresponding *vgrG1c* and *vgrG1a* double deletion (Fig. 4A-4E). Although double deletion of *tse5*/*7* and *tse6*/*7* resulted in slightly reduced T6SS assembly, introducing additional deletion of *vgrG1a* or *vgrG1c*, respectively, abolished T6SS assembly (Fig. 4A-4B, 4D and Fig. S10E-S10G). In addition, the triple effector gene deletion mutant *tse5*/*6*/*7* exhibited abolished T6SS assembly similar to the triple *vgrG1a*/*b*/*c* deletion mutant (Fig. 4A-4E). Collectively, these data indicate that the observed VgrG-mediated H1-T6SS assembly requires proper loading of VgrG-dependent effectors, which is consistent with recent findings in other species ^57,65^. Because double effector deletion mutants *tse6*/*7* and *tse5*/*7* also exhibited reduced dueling in comparison with wild type (Fig. 4A-4B, 4D), the delivered sister-cell effectors may also promote T6SS dueling.

**Fig. 4.**
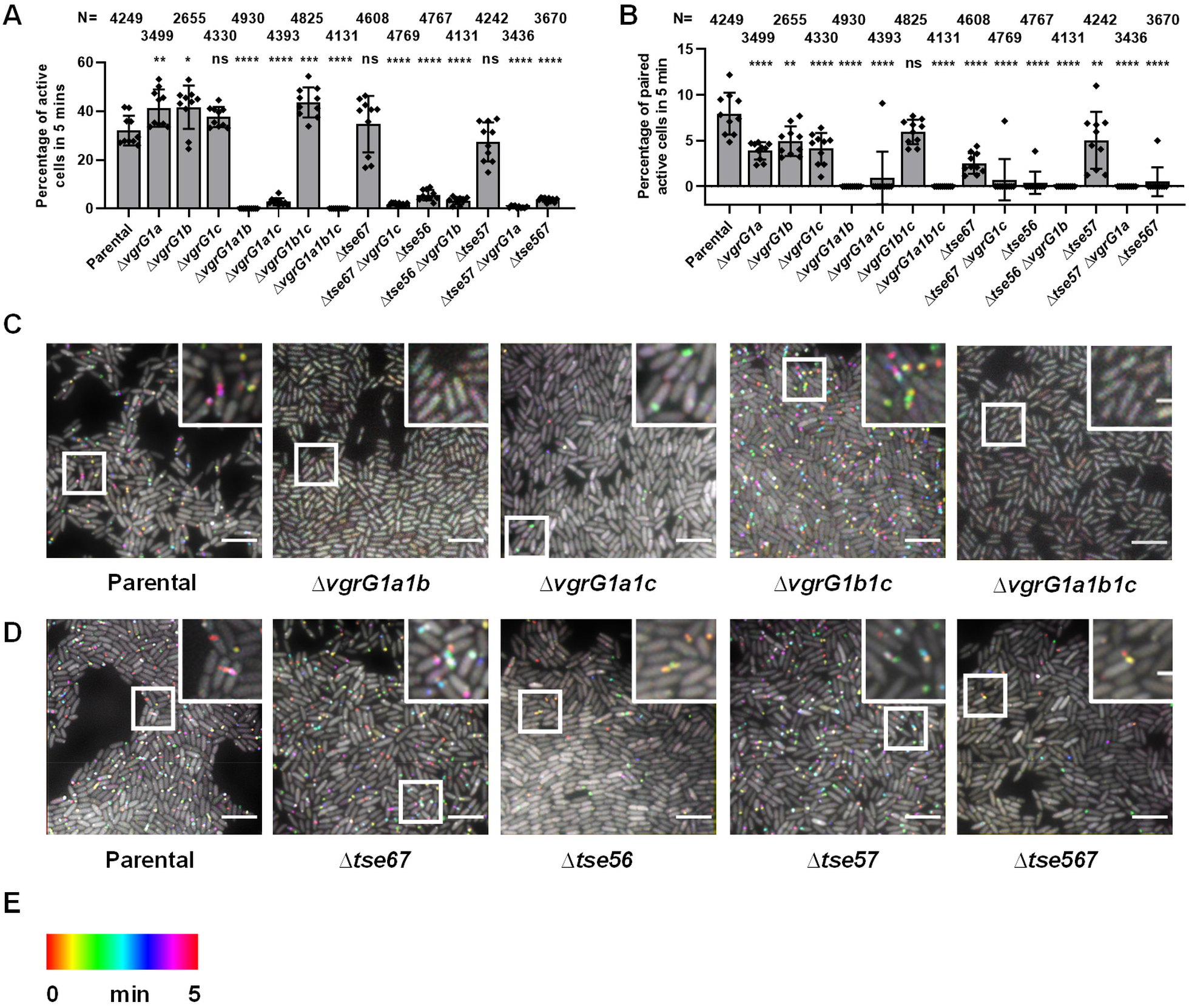
Deletion of *vgrG* and the cognate effector genes reduces T6SS dueling in *P. aeruginosa*. **A-B**, Quantification of active cells forming sheaths (A) and paired active cells (B) counted in *P. aeruginosa* H1-T6SS^+^ with combinatorial deletions of *vgrG*s and *tse5* to *tse7*. Time-lapse imaging of mutants with deletions of *vgrG* genes individually or mutants with combinatorial deletions of VgrG-dependent *tse* genes and their non-cognate *vgrG* genes is shown in Fig S10A-S10G. N indicates the total number of cells counted for each strain per a 30 μm × 30 μm field. Error bars indicate the mean ± standard deviation of at least three different biological duplicates with 10 fields. One-way ANOVA with Dunnett’s multiple comparisons test compared with the parental strain. \**P* < 0.05, \*\**P* < 0.01, \*\*\**P* < 0.001, \*\*\*\**P* < 0.0001; ns, not significant. **C-D**, Time-lapse imaging of TssB1-sfGFP signal captured every 10 s for 5 min and temporally color-coded in parental mutant H1-T6SS^+^ (Δ*retS*, Δ*tssB2*, Δ*tssB3*, TssB1-sfGFP) with the combinatorial deletions of *vgrG* genes (C) or the combinatorial deletions of VgrG-dependent *tse* genes (D). A representative image of 30 μm × 30 μm field of cells with a 2 × magnified 5 μm × 5 μm inset (marked with a box) of a selected region is shown. Scale bar in A to B is 5 μm for the large field of view and 1 μm for the insets view and applied to all. Strain genotypes are indicated at the bottom. **E**, Color scale used to temporally color code the TssB1-sfGFP signal.

### Physical contact of H1-T6SS promotes dueling

Previous studies report that all-effector-inactivated *V. cholerae* and effector-deleted *A. tumefaciens* cannot elicit a strong tit-for-tat response in *P. aeruginosa* ^5455^. To test whether the H1-T6SS dueling occurs independent of effector activities, we constructed a mutant lacking all known effectors (*8eff*_*c*_). Although the killing functions of *8eff*_*c*_ against competing *V. cholerae* and *A. dhakensis* were reduced to the T6SS-null mutant level, it exhibited active T6SS assemblies (Fig. 2A-2B, 3A-3B and Fig. S1B, S9A-S9B). Importantly, T6SS dueling was found in the *8eff*_*c*_ mutant, further supporting that effector activities of the H1-T6SS are not essential for dueling.

We recently developed a Cre-recombinase delivery system that can detect T6SS-mediated Cre delivery, which in turn causes irreversible Cre-mediated recombination changes ^36,66^. We transformed the *V. cholerae* effector mutant *4eff*_*c*_ with the pFIGR reporter plasmid carrying the two *loxP* sites flanking a gentamycin-resistance marker. We also transformed the *P. aeruginosa 8eff*_*c*_ donor with a plasmid-borne Tse6^N^-Cre^67^. Results show that the recombination efficiency in the *4eff*_*c*_ strain is comparable to the wild type and substantially higher than the Δ*tssM* mutant, indicating that the *8eff*_*c*_ could sense the physical penetration of the *V. cholerae* T6SS (Fig. 5A-5B and Fig. S11A-S11E).

**Fig. 5.**
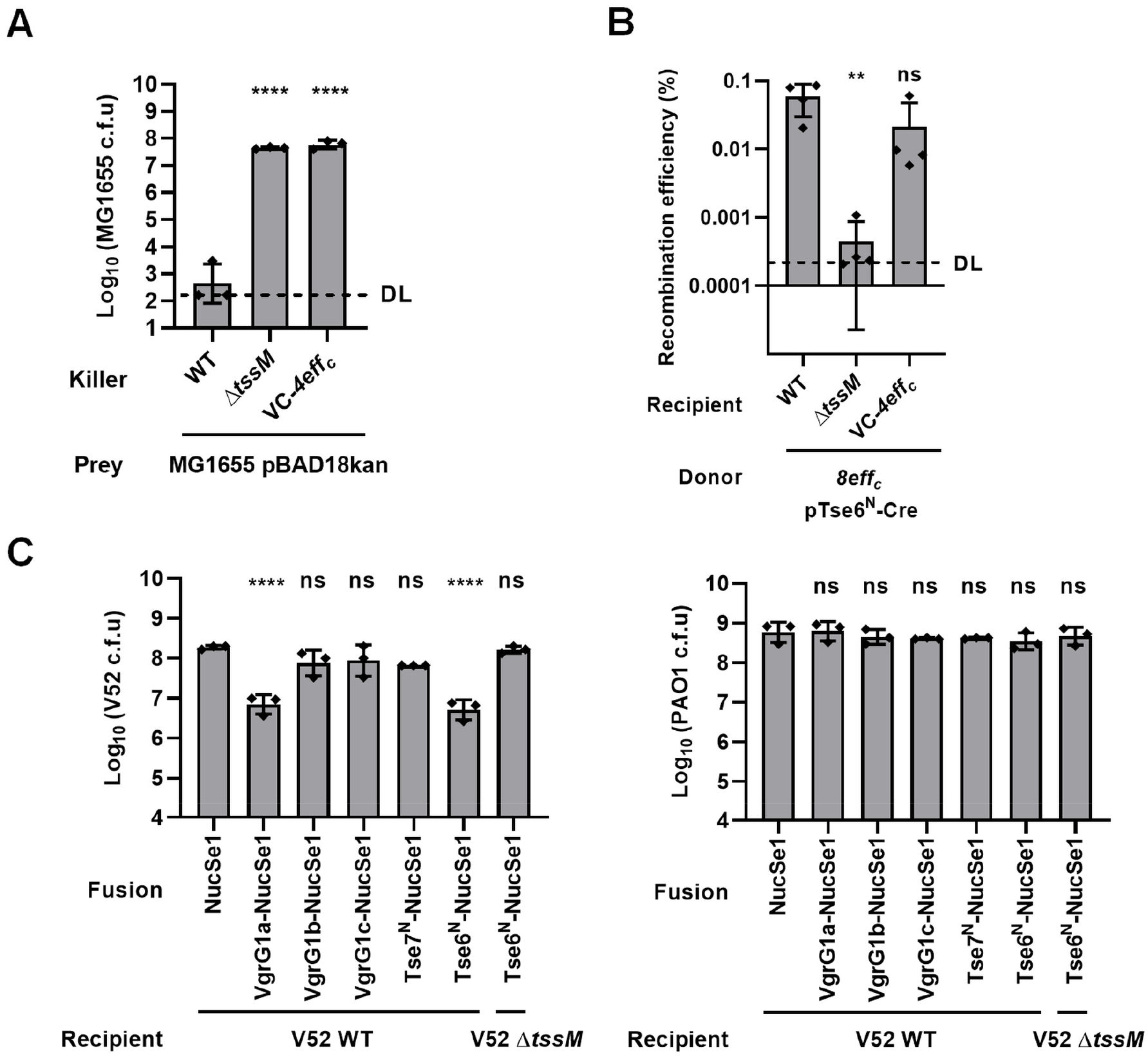
Physical contact of H1-T6SS promotes dueling. **A**, Survival of *E. coli* MG1655 with pBAD18kan plasmid after attacked by *V. cholerae* V52 wild type (WT), T6SS-null (Δ*tssM*) or VC-*4eff*_*c*_. The survival of *V. cholerae* is shown in Fig S11A. **B**, Recombination efficiency of Tse6^N^-Cre fusion delivered by *P. aeruginosa* eight-effector-inactivated mutant *8eff*_*c*_ with Tse6^N^-Cre fusion after co-incubation with *V. cholerae* WT, Δ*tssM* or VC-*4eff*_*c*_ with pFIGR plasmids. Recovery of *V. cholerae* Cre-recombined recipient cells (Gen^R^), *V. cholerae* recipient cells with pFIGR plasmid (Carb^R^), and *P. aeruginosa 8eff*_*c*_ carrying Tse6^N^-Cre fusion (Donor) is shown in Fig S11B-S11E, respectively. **C**, Survival of *V. cholerae* (left) and *P. aeruginosa* (right) after competition assay. *P. aeruginosa 8eff*_*c*_ was used control. DL, approximate detection limit. Error bars indicate the mean ± standard deviation of at least three different biological duplicates. One-way ANOVA with Dunnett’s multiple comparisons test compared with *V. cholerae* WT in A and B or compared with *P. aeruginosa* with NucSe1. \*\**P* < 0.01, \*\*\*\**P* < 0.0001; ns, not significant.

To test whether *8eff*_*c*_ could deliver other cargo proteins, we chose the NucSe1 nuclease as a cargo and constructed VgrG-NucSe1 and Tse-NucSe1 fusions^57,68^, respectively (Fig. 5C). Using *V. cholerae* strains as prey, we tested the killing efficiency of *8eff*_*c*_ carrying VgrG-NucSe1 or Tse-NucSe1 fusions and used NucSe1-only vector and *V. cholerae* Δ*tssM* prey as negative controls. Competition results show that VgrG1a-NucSe1 and Tse6^N^-NucSe1 could restore the killing activity of *P. aeruginosa 8eff*_*c*_, suggesting that NucSe1 is delivered.

## Discussion

There are mainly two modes of T6SS firing, random firing and T6SS dueling. The former has been reported in diverse T6SSs including the T6SS in *V. cholerae* and the H2-T6SS in *P. aeruginosa* ^18,69^. In contrast, dueling is a precise strategy employed by *P. aeruginosa* to selectively counterattack neighbor cells that launch the initial attacks including their sister cells, which might be more effective against a competitor than random T6SS firing. However, the molecular details of dueling remain elusive. Here, we have systematically examined the H1-T6SS dueling by constructing a series of combinatorial mutants lacking different T6SS-clusters, effectors, and VgrG proteins. Between sister cells, we show that the dueling occurs between H1-T6SS only but not between H1- and H2-T6SS. Inactivation of H1-T6SS effector functions, causing membrane and periplasmic stresses, may attenuate but cannot abolish dueling. Overexpression of Tse1, a cell-wall targeting effector, elicited strong activation of the H1-T6SS and dueling. We also reveal the differential contributions of the H1-T6SS secreted VgrG1a/b/c proteins and their associated effectors to dueling activities and T6SS assembly. The crucial role of VgrG-dependent effectors in H1-T6SS assembly further support the recent discoveries of effectors being part of a structural component ^57,65,70^. Using an all-effector-inactivated mutant, we show that effector activities are not required for the H1-T6SS. Collectively, these results suggest that both effector activities and the physical penetration of H1-T6SS contribute to the dueling activities (Fig. 6A-6C).

**Fig. 6.**
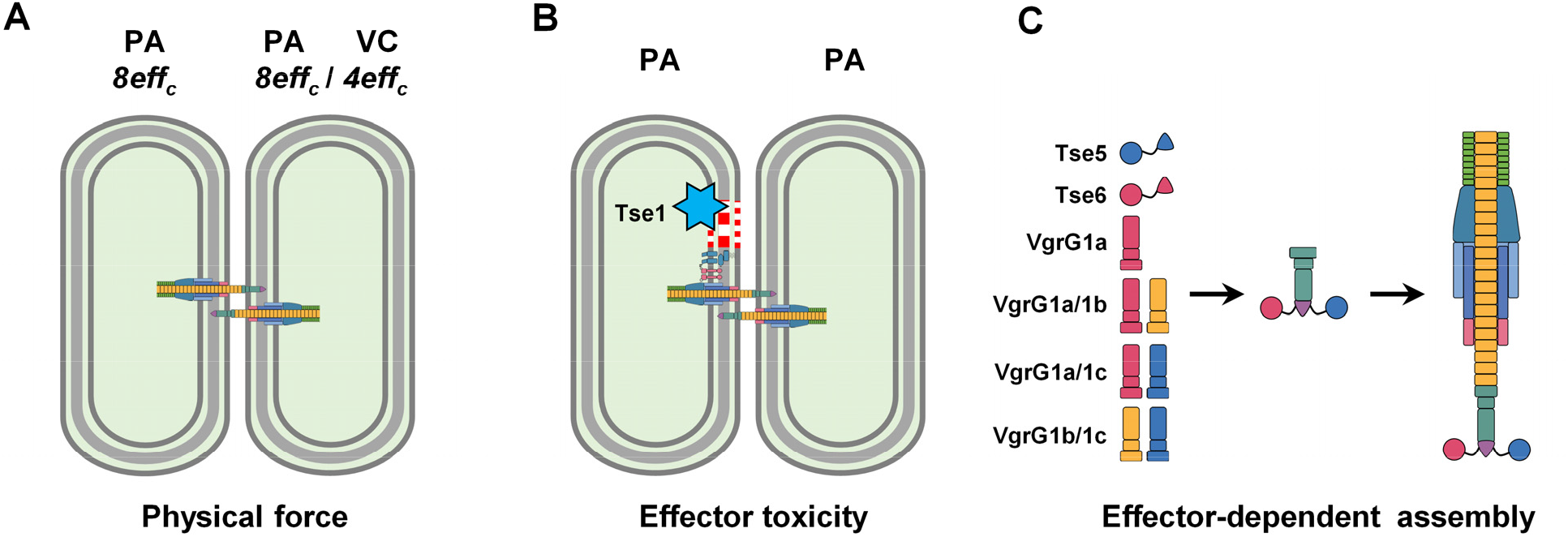
T6SS assembly and dueling is modulated by physical puncture and effectors. **A**, The T6SS physical puncture promotes dueling. **B**, The toxicity of Tse effectors targeting cell membrane or peptidoglycan could induce cell envelope disturbance, leading to the activation of TagQRST-PpkA-Fha1-PppA regulatory cascade and subsequent T6SS assembly in sister cell. **C**, H1-T6SS assembly requires proper loading of VgrG-dependent effectors Tse5 and Tse6. VgrG1a plays a key role in T6SS assembly but can be substituted by a heterotrimeric VgrG1b/c.

Activation and dueling of the H1-T6SS is controlled by the same TPP signal-transduction pathway at the post-translational level. Despite the H1-T6SS could secrete multiple toxic effectors, sister cells are protected by the cognate immunity proteins. Therefore, dueling between H1-T6SS cells would suggest that effector toxicities are not required to activate the TPP pathway. Dueling between the all-effector-inactivated *8eff*_*c*_ cells provides strong evidence supporting this. However, effectors have also been shown to promote H1-T6SS activation, such as the lipase-effector TseL in *V. cholerae* ^54^. Here we report cell-wall-lysing activities by Tse1 could also induce the H1-T6SS. The effect of effector-toxicities is consistent with the observations that both external (e.g. EDTA, eDNA, and polymyxin B) and internal membrane disturbing factors have been shown to activate the H1-T6SS through the TPP pathway ^47,50,56^. Therefore, the TPP-mediated activation is likely dependent on membrane-perturbing signals but does not differentiate whether the signals are from physical penetration or effector-toxicities.

Analogous to a power drill, the spear-like T6SS relies on an instant and strong power to pierce through the cell envelope of both gram-negative and gram-positive cells, as evidenced by the cytosolic delivery of structural components and nuclease and other cytosolic toxins ^68,69,71–75^. Considering the frequent T6SS-firing between sister cells, the physical force is obviously insufficient to harm bacteria. However, the puncture from the all-effector-inactivated detoxified T6SS in *V. cholerae* increases membrane permeability and reduces survival of *E. coli* envelope-impaired mutants, suggesting an imposed membrane stress by the incoming T6SS spear-like tube and its physical force ^54^. How cells respond to such force and repair damages remains unclear and may vary among different species. For the interaction between *V. cholerae* and *P. aeruginosa*, the all-effector-inactivated *V. cholerae* T6SS elicits a much weaker dueling/tit-for-tat response from *P. aeruginosa* during a competition assay, in contrast to the strong response induced by the *V. cholerae* lipase effector TseL ^54^. This led us to propose that effector functions but not the physical force of the T6SS are the primary trigger for the tit-for-tat response ^54^. However, the competition assay is prone to be disturbed by environmental factors, including divalent cations and temperature ^66^, and is less sensitive to the Cre-delivery, which causes irreversible genetic changes ^36,66^. Using a Cre-dependent reporter, we report that the *V. cholerae* T6SS physical contact could also elicit a dueling response. Interestingly, the H2-T6SS has little effect on inducing H1-T6SS. How can the H1-T6SS differentiate the H1-T6SS attack from the H2-T6SS attack? It is possible that the H2-T6SS activation condition may be repressive to the H1-T6SS, or that the H1-T6SS is more sensitive to itself due to the recyclable structural components, VgrGs, and effectors, which promote T6SS assembly when delivered from a sister cell. Nonetheless, these data suggest the H1-T6SS can be activated by at least some types of physical puncture.

Finally, the detailed dissection of effector functions is important to reveal the differential contribution of effectors to the T6SS assembly and antibacterial activities toward specific pathogens. For example, of the H1-T6SS eight effectors, Tse5 and Tse6 directly contribute to assembly as structural components since lacking both severely impaired T6SS functions. Further research is required to elucidate the molecular details of unique defense pathways accounting for the different sensitivities of these pathogens. These findings may not only help better understand how the T6SS is assembled and evolved in the acquisition of multiple effectors but also reveal novel antibiotic targets to inhibit diverse defense systems and to treat complex polymicrobial infections.

## Materials and Methods

### Bacterial strains and growth conditions

Strains, plasmids, and primers used in this study are listed in Table S1. Cultures were grown in LB (1% [w/v] tryptone, 0.5% [w/v] yeast extract, 0.5% [w/v] NaCl) aerobically at 37 °C. Antibiotics were used at the following concentrations: streptomycin (100 µg/ml), irgasan (25 µg/ml), gentamicin (20 µg/ml), and kanamycin (50 μg/ml).

### Construction of mutants

Homologous recombination and suicide vector pEXG2.0 were used to construct all mutants as previously described ^30^. Amplified homologous arms were overlapped by PCR and inserted into pEXG2.0 by Gibson assembly. The pEXG2.0 plasmids carrying homologous arms were transformed into donor *E. coli* SM10 λ pir or WM6026. The donor and recipient strain were mixed and incubated at 37 °C for about 3 h. Trans-conjugants were screened on LB plates with selective antibiotics. Plates with 6% sucrose were used to select sucrose-resistant colonies at 22 °C. All mutants were confirmed by PCR or Sanger sequencing.

### Construction of expression plasmids

For expression vectors, genes of interest were amplified by PCR and cloned into pPSV37 plasmids using Gibson assembly. All constructs were confirmed by PCR and Sanger sequencing. All plasmids and primers used in this study are listed in Table S1.

### Bacterial cell killing assay

Single colonies of killer cells were cultured overnight in LB with appropriate antibiotics, and 1/50 (for *P. aeruginosa)* or 1/100 (for *V. cholerae*) sub-cultured to OD_600_∼1. Killer cells were collected at 10, 000 × *g* for 30 seconds twice, and resuspended in 100 µl LB to OD_600_=10. For competition assay of *P. aeruginosa* against different T6SS^+^ prey cells, overnight cultures of prey cells on LB-agar plates were normalized to OD_600_=2 and mixed with killer cells at a ratio of 5: 1 (killer: prey). For competition assay between *V. cholerae* and *E. coli* MG1655 with pBAD18kan plasmid, overnight cultures of MG1655 were normalized to OD_600_=1 and mixed with *V. cholerae* at the ratio of 10: 1(killer: prey). For competition assay of *P. aeruginosa* with NucSe1 fusions against *V. cholerae*, overnight cultures of *V. cholerae* were normalized to OD_600_=1 and mixed with *P. aeruginosa* carrying NucSe1 fusions at the ratio of 10: 1 (killer: prey). IPTG (isopropyl-β-D-thiogalactoside, 1mM) was used to induce the expression of NucSe1 fusions. Mixtures were incubated at 37 °C for 3 h and resuspended in 500 μl fresh LB. A series of 10-fold serial dilutions were performed and LB-agar with the appropriate antibiotic were used to assess the survival of killer and prey cells. LB-agar plates containing 100 μg/ml streptomycin were used to select for *V. cholerae* V52, *A. dhakensis* SSU, LB-agar plates with 25 μg/ml irgasan were used to select *P. aeruginosa*, and LB-agar plates with 50 μg/ml kanamycin were used to select *E. coli* MG1655 with pBAD18kan. All competition assays were performed at least triplicate. The mean Log_10_ c.f.u of recovered prey or killer were plotted, and error bars show the mean ± standard deviation of three independent biological replicates. 0.5 colonies were counted if no colonies were observed and labeled as the detection limit (DL) as previously described ^76^. One-way ANOVA with Dunnett’s multiple comparisons test or a two-tailed Student’s *t*-test was performed to determine *P*-values.

### Cre delivery assay

Single colonies of *P. aeruginosa* cells were grown in 500 µl LB with appropriate antibiotics and 1/50 sub-cultured to OD_600_ ∼1. *P. aeruginosa* cells were collected at 10, 000 × *g* for 30 seconds twice, resuspended in fresh LB to OD_600_ =10. Overnight cultures of recipient cells (*V. cholerae* strains with pFIGR) were normalized to OD_600_ =1. Donor and recipient cells were mixed at a ratio of 10: 1. LB-agar plates with 1 mM IPTG were used to induce the expression of Cre fusions. After co-incubation for 3 h at 37 °C, the mixtures were resuspended in 500 μl fresh LB and a series of 10-fold serial dilutions were performed. LB-agar plates with the appropriate antibiotic were used to assess the survival of donor and recipient cells. LB-agar plates containing 100 μg/ml streptomycin and 20 μg/ml gentamicin were used to select for *V. cholerae* with obtained gentamicin resistance, 100 μg/ml streptomycin and 100 μg/ml carbenicillin for original *V. cholerae* recipient cells, 100 μg/ml streptomycin for recovery of *V. cholerae* cells, and 25 μg/ml irgasan and 20 μg/ml gentamicin for the recovery of *P. aeruginosa* cells. Data are shown as Log_10_ c.f.u for survival of donor or recipient cells on LB plates with selective antibiotics or recombination efficiency, calculated as Gen^R^ c.f.u/Carb^R^ c.f.u. If no colonies were observed, samples were listed as the detection limit (DL), shown as if 0.5 colonies were counted ^76^. Error bars show the mean ± standard deviation of three biological replicates. One-way ANOVA with Dunnett’s multiple comparisons test was used to determine *P*-values.

### Fluorescence microscopy image acquisition and analysis

Single colonies of *P. aeruginosa* were grown in 500 µl LB overnight and 1/50 sub-cultured in 3 ml LB with 20 μg/ml gentamicin and 25 μg/ml irgasan for *P. aeruginosa* carrying pPSV37 expression vectors or 25 μg/ml irgasan for *P. aeruginosa* mutants. Cells were grown to OD_600_∼0.8-1, and 1 mM IPTG was used to induce the expression of pPSV37 plasmids for about 1 h. Cells were spotted on 1% agarose-0.5 × PBS pads. Regions on the edges were taken for T6SS dueling while random areas were taken when Tse effectors were expressed in PAO1 strains. All the microscopy images were obtained using a Nikon Ti-E inverted microscope with a Perfect Focus System (PFS) and a CFI Plan Apochromat Lambda 100 × oil objective lens. ET-GFP (Chroma 49002) filter sets were used for the GFP fluorescence signal and ET-mCherry (Chroma 49008) filter sets were used for the mCherry2 fluorescence signal.

Fiji software was used to analyzed all images. Images from a 5 mins series, 10 s interval were normalized to the same mean intensity to correct for photobleaching, as described previously ^77^. Total cell numbers were calculated from the phase-contrast figure, adjusting the threshold to demarcate the boundary of cells and using the “analyze particles” option, excluding particles on the edge. Cell counts were confirmed manually. To ensure the accuracy and repeatability of the results, at least 6 fields were picked for quantification of active cells and paired active cells. The plugin “Temporal-Color code” for Fiji was used to assess the active sheath over the time frame. Positive sfGFP or mCherry2 cells were counted manually. Contrast within compared image sets was adjusted equally. All imaging experiments were performed with at least three biological replicates.

### Tse effector toxicity assay

Cells expressing different plasmid constructs were streaked onto LB-agar plates supplemented with 0.2% [w/v] glucose at 37 °C overnight. Cells were then collected and resuspended in fresh LB and normalized to OD_600_=1. A series of 10-fold dilutions were spotted on LB-agar plates with 0.2% [w/v] glucose or 1 mM IPTG (for pPSV vectors). Each experiment was repeated three times, with one representative experiment shown.

### Statistical analysis

Graphpad Prism software (8.2.1) was used to analyze all data. One-way ANOVA with Dunnett’s multiple comparisons test or two-tailed Student’s *t*-test was used to calculate *P*-values as indicated.

## Data availability

The source data underlying Figs. 1C-1D, 2A-2D, 3A-3C, 4A-4B, 5A-5C and Figs. S7I, S9A-S9C, S11A-S11D are provided as a Source Data file. Other data supporting the findings of this study are available within the paper or from the corresponding author upon request.

## Acknowledgments

This work was supported by funding from the National Key R&D Program of China (2020YFA0907200), National Natural Science Foundation of China (32030001), and National Natural Science Foundation of China-Youth Science Fund (82102399). We thank Kevin Manera and Steve Hersch for their helpful comments. The funders had no role in study design, data collection and interpretation, or the decision to publish.

## Author contributions

T.D. conceived the project. L.W. performed most experiments, analyzed results and prepared all figures. T.P. contributed to plasmid construction; L.W., M.T., L.W., and S.Y. contributed to competition assay. H. L. contributed to strain construction. X.L. contributed to the experimental design. L.W., S.S., X.L., and T. D. wrote the paper.

